# Predation and competition drive trait diversity across space and time

**DOI:** 10.1101/2023.04.05.535749

**Authors:** Zoey Neale, Volker H. W. Rudolf

**Affiliations:** Graduate Program in Ecology & Evolutionary Biology, BioSciences Rice University, Houston, TX 77005, USA

**Keywords:** Functional diversity, beta diversity, Odonata, meta-community, seasonal, spatial, aquatic

## Abstract

Competition should play a key role in shaping community assembly and thereby local and regional biodiversity patterns. However, identifying its relative importance and effects in natural communities is challenging because theory suggests that competition can lead to different and even opposing patterns depending on the underlying mechanisms. Here, we’ve taken a different approach: rather than attempting to indirectly infer competition from diversity patterns, we compared trait diversity patterns in odonate (dragonfly and damselfly) communities across different spatial and temporal scales along a natural competition-predation gradient. At the local scale (within a community), we found that trait diversity increased with the size of top predators (from invertebrates to fish). This relationship is consistent with differences in taxonomic diversity, suggesting that competition reduces local trait diversity through competitive exclusion. Spatial (across communities) and temporal (within communities over time) trait variation peaked in communities with intermediated predators indicating that both high levels of competition or predation select for trait-convergence of communities. This indicates that competition acts as a deterministic force that reduces trait diversity at the local, regional, and temporal scales, which contrasts with patterns at the taxonomic level. Overall, results from this natural experiment reveal how competition and predation interact to shape biodiversity patterns in natural communities across spatial and temporal scales and provide new insights into the underlying mechanisms.

**Open Research statement:** Data and/or code are provided as private-for-peer review via the following link: [Link to external storage location] and will be made publicly available with publication

## Introduction

The number and distribution of species present in a given habitat typically vary even among close neighbors within a geographic region. To help explain this variation, community assembly theory has traditionally focused on the concept of “filters” for describing the processes that determine which species may exist within a local habitat (Keddy 1992, Belyea and Lancaster 1999, Chase 2003, Götzenberger et al. 2012, HilleRisLambers et al. 2012). The dispersal “filter” restricts the available regional species pool (i.e. which species can reach a local site), while the abiotic filter selects for species with certain traits that allow them to persist under the environmental conditions of the local habitat. Finally, species must pass through a biotic filter, where species interactions like predation and competition limit which of these species persist (HilleRisLambers et al. 2012, Kraft et al. 2015). While this filtering framework has been broadly applied to help describe the key processes that drive variation in community composition (Keddy 1992, HilleRisLambers et al. 2012, Kraft et al. 2015, Germain et al. 2018), it relies on the assumption that different filters leave clear signals, especially in trait (functional) diversity patterns. However, the expected signal of some biotic filters (e.g. competition) remains heavily debated with seemingly contradicting predictions making it difficult to identify the relative importance of different underlying processes (Mayfield and Levine 2010). As a consequence, it remains unclear when and how biotic processes like competition affect biodiversity patterns, especially at regional scales.

Identifying the relative importance of biotic factors in driving biodiversity patterns is challenging because different patterns can arise from the same processes (Mayfield and Levine 2010), and multiple processes can interact with each other (Dunson and Travis 1991). For instance, competition is clearly a key factor determining species coexistence within a given community and thus should act as a “filter” that determines which species are present (Chesson 2000, Friedman et al. 2017). But, coexistence theory (Chesson 2000) suggests that the resulting trait diversity patterns can differ substantially depending on which processes are operating. On one hand, asymmetric (hierarchical) competition for a shared limiting resource may select for an optimal suite of traits that maximize a competitor’s ability to acquire the resource. This would lead to a negative relationship between competition strength and functional diversity (Ackerly and Cornwell 2007). On the other hand, the theory of limiting similarity posits that competing species must demonstrate sufficient differences in their niches to persist, which would lead to a positive relationship between competition strength and functional diversity (MacArthur and Levins 1967, Meszéna et al. 2006). Thus, competition could either enhance or reduce functional diversity within a given community (Mayfield and Levine 2010). However, strong abiotic filters can also shape (increase or decrease) functional diversity, making it challenging to isolate and infer how competitive processes shape functional diversity patterns within communities from purely observational data (Dunson and Travis 1991, Mayfield and Levine 2010).

Much less is known about how competition affects biodiversity patterns at the regional scale or the temporal scale, i.e. how functional diversity varies across sites within a region (spatial β-diversity) or within a community over time (temporal β-diversity). Regional diversity patterns are often explained by examining the relative strength of stochastic vs. deterministic processes (Chase 2007, Chase et al. 2009, Van Allen et al. 2017). Stochastic processes are expected to increase variation in species composition across habitats and within habitats over time, and thus positively affect spatial beta diversity (Chase 2007, Chase et al. 2009). In contrast, deterministic processes should reduce variation in community composition across habitats and thus reduce spatial β-diversity with more predictable changes in compositions over time (i.e. similar seasonal composition across years) (Amarasekare 2003, Chase 2003, 2007). Competition and priority effects are often proposed as a stochastic process that promotes high β-diversity at the species level (Fukami 2004, Chase 2007). But this may not hold at the trait level where it should depend on the specific type of competition. If species compete for a limited set of resources (niches), we expect the same trait combinations in similar habitats and over time within communities. In this scenario, competition would act as a deterministic force on trait diversity. In this scenario, spatial β-diversity might still be high at the species level if the species pool includes functionally redundant species (i.e. species with similar trait combinations) and arrival times vary across time and space. Thus, analyzing both local and regional trait diversity patterns and comparing them to species diversity patterns may help to provide more nuanced insights into how important and what type of competition is in shaping biodiversity patterns.

Capturing competition strength gradients in a field setting can be difficult, as manipulative experiments are often needed to directly measure the strength of competition. However, most systems demonstrate an inverse relationship between competition and predation strengths (Paine 1966, Morin 1981, 1983, Sih et al. 1985, Gurevitch et al. 2000, Chase et al. 2002, Terborgh 2015, Ellingsen et al. 2020). This provides the ability to infer competition strength from predation strength in systems where predation strength is observable and the inverse relationship between competition and predation has been demonstrated. However, in these scenarios, predictions must be modified to account for the effects of predation strength on functional diversity. If greater competition leads to higher levels of functional diversity (limiting similarity), we expect to see a monotonic decline in functional α-diversity as competition strength decreases with increased predation. However, if competition leads to increased clustering of traits, we expect the opposite trend along the predation strength gradient. At the regional scale, previous studies indicate that β-diversity decreases with predation at the species level (Chase et al. 2009, Van Allen et al. 2017). If competition acts as a stochastic driver then we expect the same relationship for trait β-diversity. However, if competition acts as a deterministic force, functional β-diversity should not decline and instead peak at intermediate levels of predation.

To test these alternate predictions, we analyzed the spatiotemporal patterns of functional diversity in habitats spanning a natural competition-predation gradient. We used a time series of trait diversity patterns of in odonate (dragonfly and damselfly) larvae in natural ponds that differ in their top predators to answer the following questions: (1) How does local (alpha) trait diversity change in response to the relative strength of competition vs. predation. (2) Are there consistent differences in spatial and temporal beta trait-diversity patterns across this predation gradient? (3) Finally, we ask how similar patterns in trait diversity are to species diversity patterns in this system.

## Methods

### Study System

We conducted this study on larval odonate communities in freshwater ponds in the Piney Woods region of eastern Texas. The ponds are located in Davy Crockett National Forest (n = 36) and Angelina National Forest (n = 9). Larval odonates show a high taxonomic (>36 species) and trait diversity in this region and presence and abundance vary considerably across species, sites, and within sites over time (Van Allen et al. 2017). The study ponds varied widely in area (564 – 1393m^2)^, depth, and level of permanence and fall along a natural gradient of predation strength. The top predators were, from weakest to strongest predation strength on odonates, invertebrates, salamanders, green sunfish, and largemouth bass (Van Allen et al. 2017). The identity of top predators plays a key role in determining the species composition of odonate communities and strength of competition (Wellborn et al. 1996, McPeek 1998) and is by far the single best predictor of variation in odonate species composition across these ponds (Van Allen et al. 2017).

### Sample Collection

We collected seasonal community samples four times per year (spring, summer, fall, and winter) beginning in the summer of 2008 and ending in the fall of 2011 (n = 14 samples per pond). We used dip nets to collect specimens that we preserved in alcohol and brought them back to the lab for identification. Each sampling effort was approximately 24 person-minutes (22.22 ± 7.64 *SD* person-minutes). Four ponds were sampled separately for 60 person-minutes in 2011 to determine if the 24-person-minute method was an accurate sample. We found that the 24-person-minute samples collected approximately 86% (± 10% SD) of the species richness collected in 60 person-minutes.

A total of 18,891 specimens spanning 36 taxa (see **Appendix S1: Table S1** for the complete species list) were collected during the duration of this study. Most were identified to the species level, but specimens of the genus *Libellula* were identified to genus level because the species in the area could not be accurately distinguished.

### Pond habitat characteristics

We conducted predator surveys four times per year from 2007-2010 (n = 13) using similar methods as the odonate surveys described above, but predator surveys lasted 60 person minutes and predators were released. These surveys were supplemented with hook and line surveys to confirm whether fish predators were present. Any predator detected in at least two predator surveys was considered a member of the community, and predators with the largest body size were considered the top predators. Additionally, we measured pond area and depth and rated the amounts of aquatic vegetation and canopy cover as indexes from 0 to 4 to include as covariates in our models, as we believed they may improve model fit.

### Functional trait measurements

We collected twelve trait measurements (Table 1) per species. We chose these traits based on previous work in this taxa to capture differences in habitat use, predator avoidance, food resource acquisition, and mobility (Wellborn et al. 1996). Traits were measured in at least 3 randomly selected F-0 instars per species using ImageJ (Software version) and values were averaged by species. We excluded any species for which a minimum of 3 F-0 instars were not collected from analyses to avoid spurious trait measurements from biasing results. These species were rare in these communities and likely transient. This resulted in total of 27 species with trait data that were used in the final analysis.

**Table 1:**
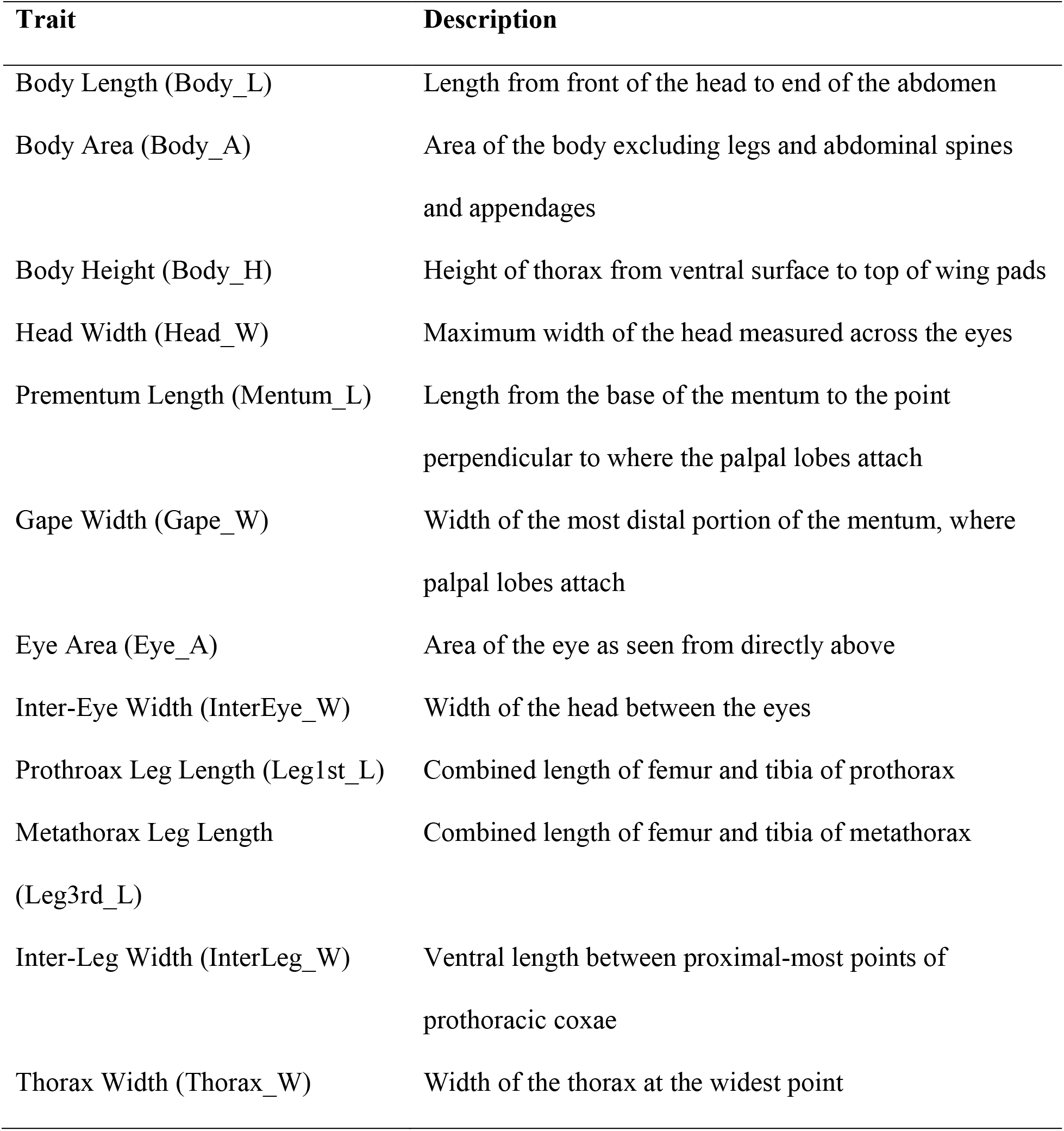
Odonate functional traits used to study trait diversity patterns. These traits are correlated with various functions, such as diet, micro-habitat use, behavior, trophic position, and vulnerability to different top predators.

### Local diversity calculation and analyses

We reduced the twelve traits to a single functional dispersion metric (FDis) for a more intuitive analysis of the functional diversity. This metric is commonly used in functional diversity studies because it is independent of species richness, can be weighted by abundance, and is insensitive to outliers (Laliberté and Legendre 2010). Briefly, to calculate FDis species traits are reduced to the first two principal components of analysis (PCoA) axes to calculate the abundance-weighted average distance of species present to a community’s centroid (**Fig. 1** for positioning of species in the multidimensional trait space). We estimated FDis for each sample as a measure of functional α diversity using the R package “FD”. Higher values of FDis indicate a greater variation of traits within ponds and thus greater functional diversity.

**Figure 1:**
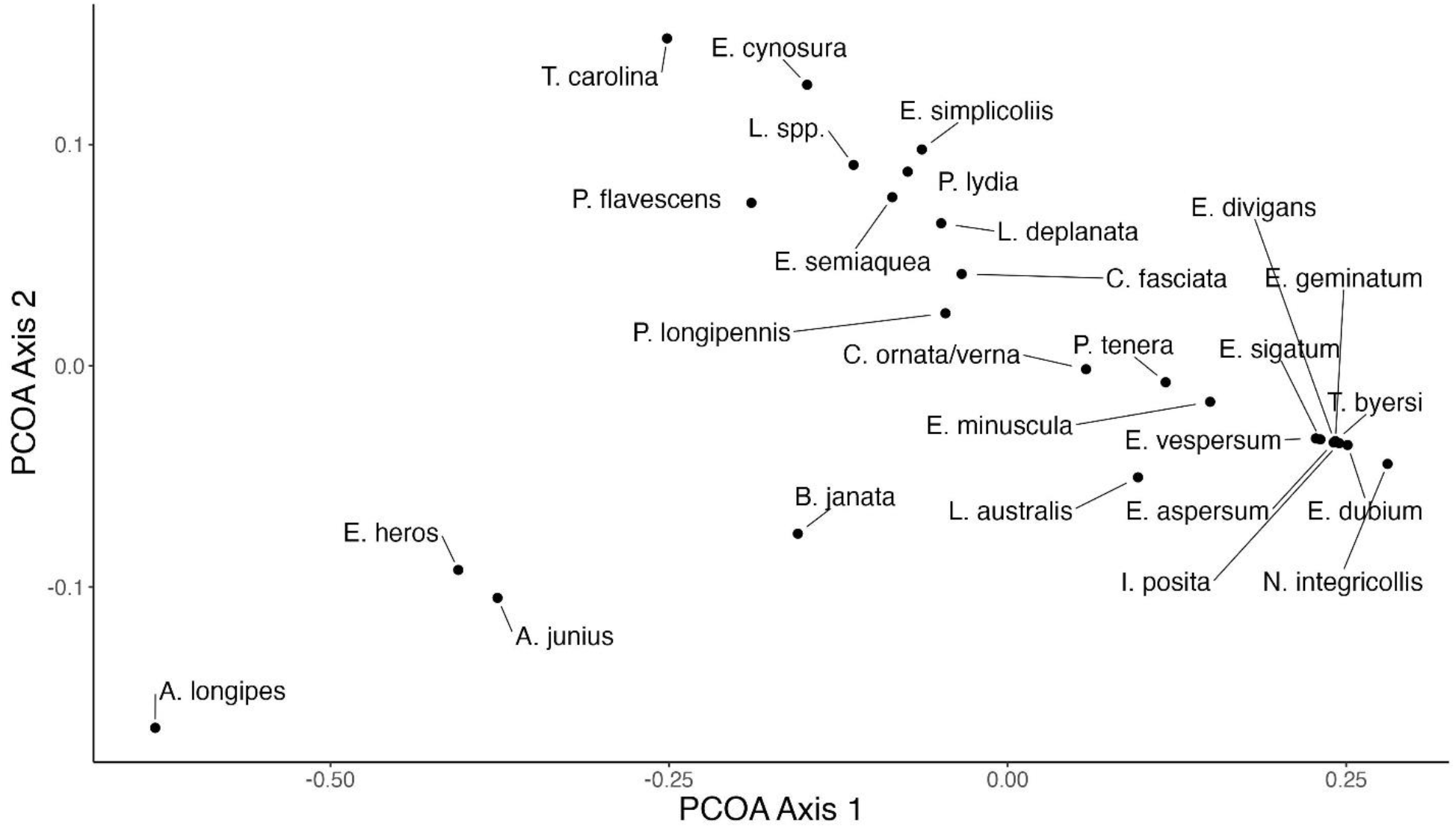
Trait diversity and differences in the odonate meta-community. Each point represents the position of a species in multidimensional trait space based on 12 traits in Table 1.

We used a generalized linear mixed model to test the fixed effects of top predator, year, season, both two-way interactions, and the random effect of pond identity on sample FDis values. Additionally, linear models including all possible combinations of pond area, depth, vegetation index, and canopy cover were tested to maximize the model fit. However, none of these pond traits improved the fit (based on AICc values) of any models including top predator and thus were excluded from the final analyses presented here (Table S2. All GLMM’s were conducted with the “lmer” function in the R package “lme4”).

### Regional diversity and analyses

We calculated differences in trait composition between ponds as measures of regional (β) functional diversity using all-pairwise Gower’s dissimilarities (Gower 1971) based on average trait values. Samples that included fewer than three individual odonates were not included.

Spatial β diversity values were extracted as the subset of pairwise dissimilarities between all ponds within the same top predator classification at the same year × season combination. In addition, we also calculated temporal β diversities to quantify changes in trait composition over time. Our previous study indicates that ponds with different predators systematically differ in how species diversity is partitioned across time and space (Van Allen et al. 2017). Thus, we also quantified temporal β diversity as the trait dissimilarities between each pond and itself at the following time step (*e.g.,* the dissimilarity between one pond in spring and the same pond in summer of the same year). Higher values indicate a greater change in species traits between seasons. Together, these analyses allowed us to test if and how the strength of stochastic vs. deterministic processes differed between ponds along the competition-predation gradient.

We used generalized linear mixed models with gamma-distributed error models and inverse link functions to test the fixed effects of top predator, year, season, and both two-way interactions with top predator on spatial and temporal β diversities, separately. Pond identity was included as a random effect in the temporal β diversity analysis and identities of both ponds compared in each dissimilarity were included as separate random effects in the spatial β diversity analysis. The season and year of the earlier sample were used in the temporal β diversity analysis. A value of 10^-5^ was added to all dissimilarity values to eliminate zeroes and allow fitting of the gamma error distribution. We conducted analyses using the “glmer” function with optimizers “optimx” and “bobyqa” in the R package “lme4.”

### Accounting for unequal sample sizes

We used a randomization test to examine the influence of unequal sample sizes in pond types on diversity metrics. This was done by randomly selecting five ponds from salamander, sunfish, and bass predator types and average α (FDis), spatial β (Gower’s dissimilarity), and temporal β (Gower’s dissimilarity) values were calculated. Five was chosen as the number of ponds to resample because this was the sample size for invertebrate ponds in the study, the predator category with the fewest number of ponds. This process was repeated 500 times for each metric, and the distribution of resampled mean diversity values was compared to the observed values.

## Results

### Trait differences across habitats

The trait combination within a community varied with predator type and season (**Fig. 2, 3**). On average, ponds with green sunfish and largemouth bass had very similar mean trait combinations (similar centroids in **Fig. 2**) and showed similar seasonal shifts. In both community types, mean trait combinations differed across seasons, but annual variation within seasons largely followed along the same diagonal axis in the nMDS space (**Fig. 3**). Trait combinations in ponds with were intermediate showing overlap in trait space between fish and invertebrate ponds depending on the specific season. Traits in invertebrate ponds were most different from fish ponds (**Fig. 2**). On average odonates were longer in salamander and invertebrate ponds, but this varied across seasons with the largest difference in winter and fall (**Appendix S1: Fig. S1 -S4**). Overall, these patterns confirm that habitats with different types of top predators select for different traits in odonates.

**Figure 2:**
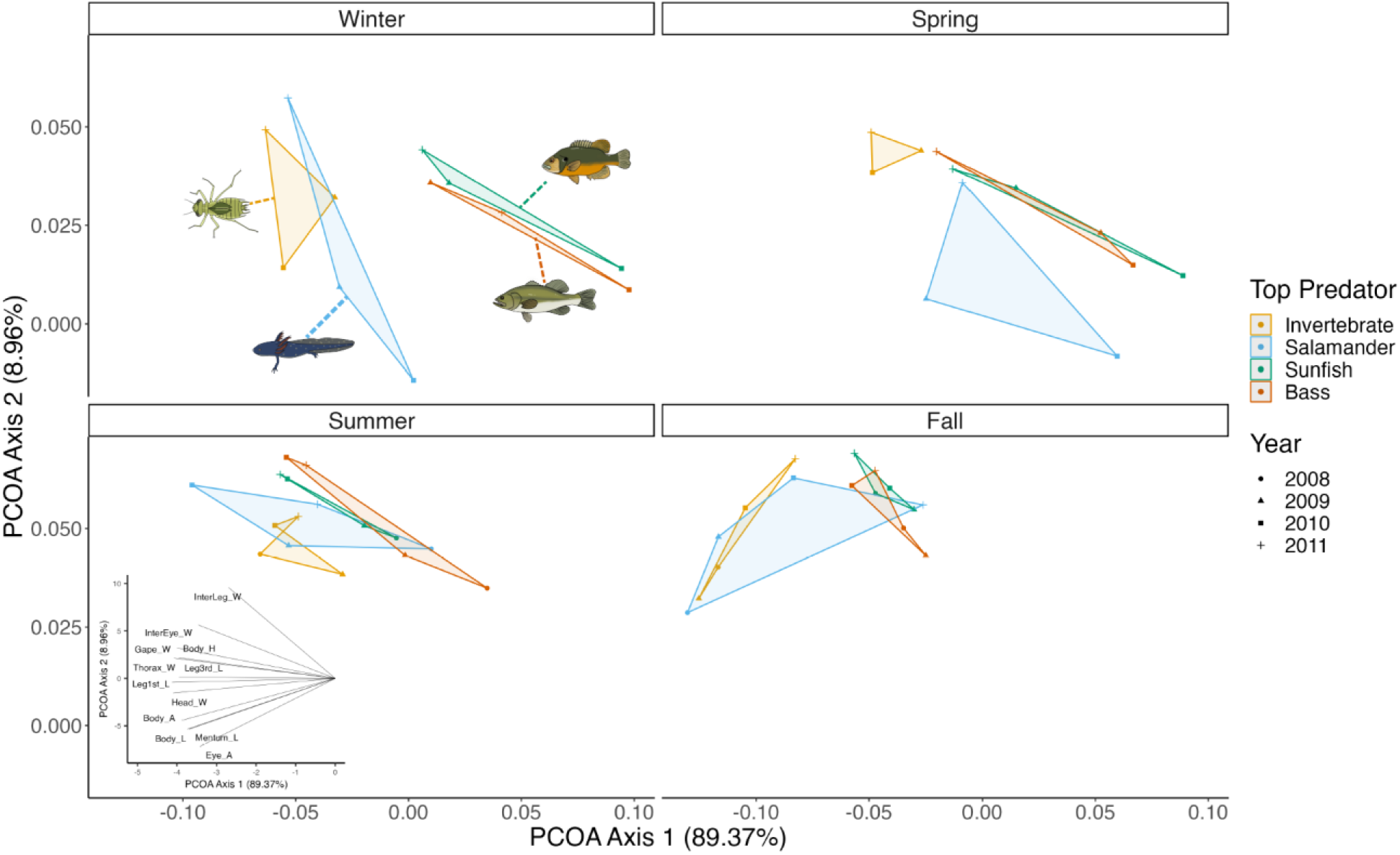
Annual and seasonal shifts in trait composition in habitats with different top predators. Points indicate average pond-level trait centroids on PCOA axes, colors indicate seasons, and shapes indicate the year of observation. Shaded areas of convex hulls indicate inter-annual variation in trait compositions.

**Figure 3:**
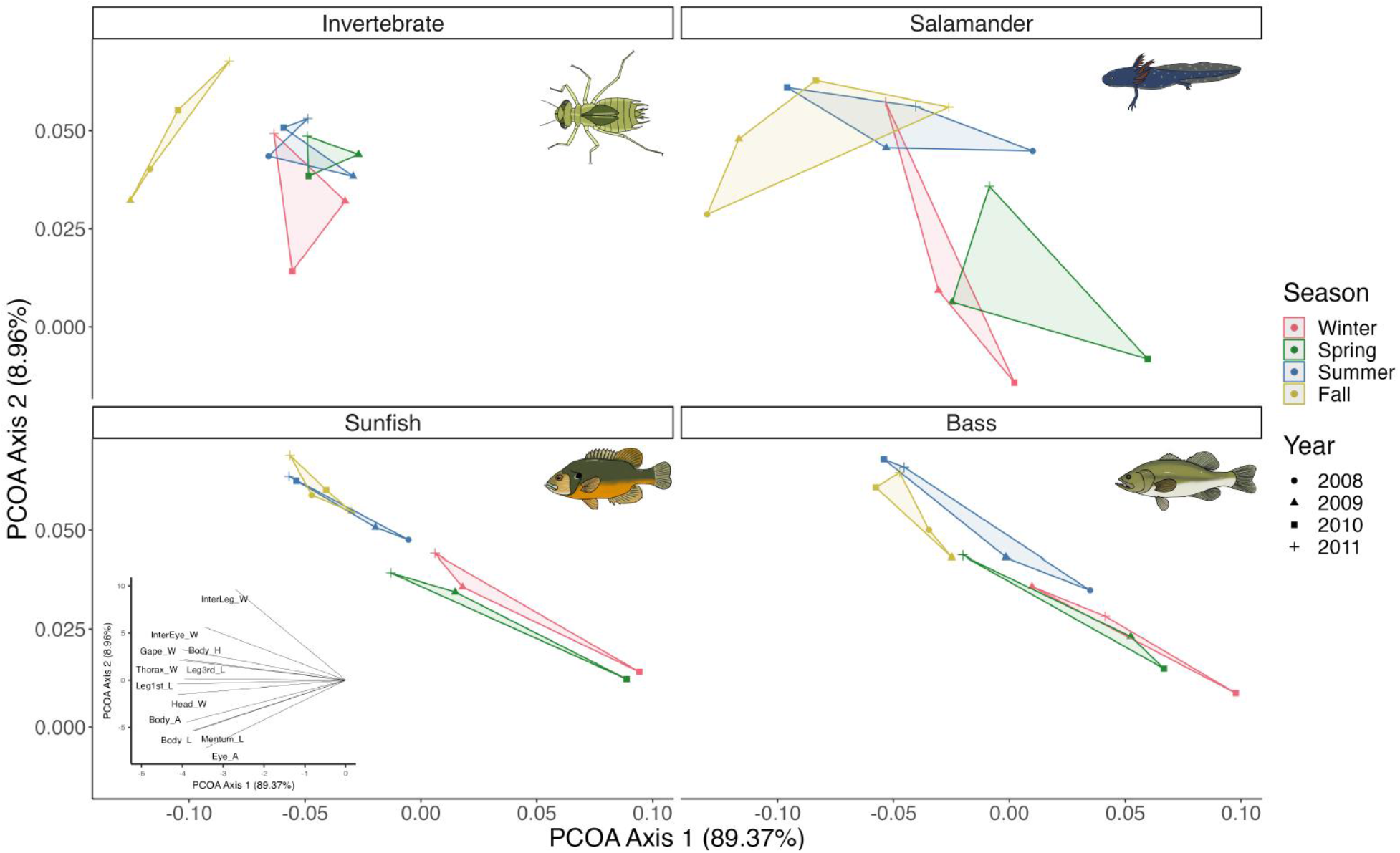
Differences in trait composition across habitats with different predators by seasons. Colors indicate the top predator in a habitat and shapes indicate the year of observation. The shaded area of the convex hull indicates interannual variation in trait compositions.

### Local (α) trait diversity

Local (within pond) trait diversity (FDis) differed across systems with different top predators, but this effect was season-specific (predator*season: χ^2^ = 19.77, p = 0.019).

Functional dispersion was typically highest in salamander ponds, except in the summer when bass ponds displayed the greatest values (**Fig. 4a**). The largest difference between top predators occurred in the spring, with salamander ponds displaying greater FDis than invertebrate ponds. The distribution of randomized values simulating equal sample sizes was similar to the observed distribution (**Appendix S1: Fig. S5**) indicating that these results were not driven by predator identity nor by sample size differences. Diversity also showed a general seasonal pattern (χ^2^ = 19.57, p = 0.002) with the highest trait diversity during winter and lowest during spring for invertebrate ponds and during summer or fall for other pond types (**Fig. 4a**).

**Figure 4.**
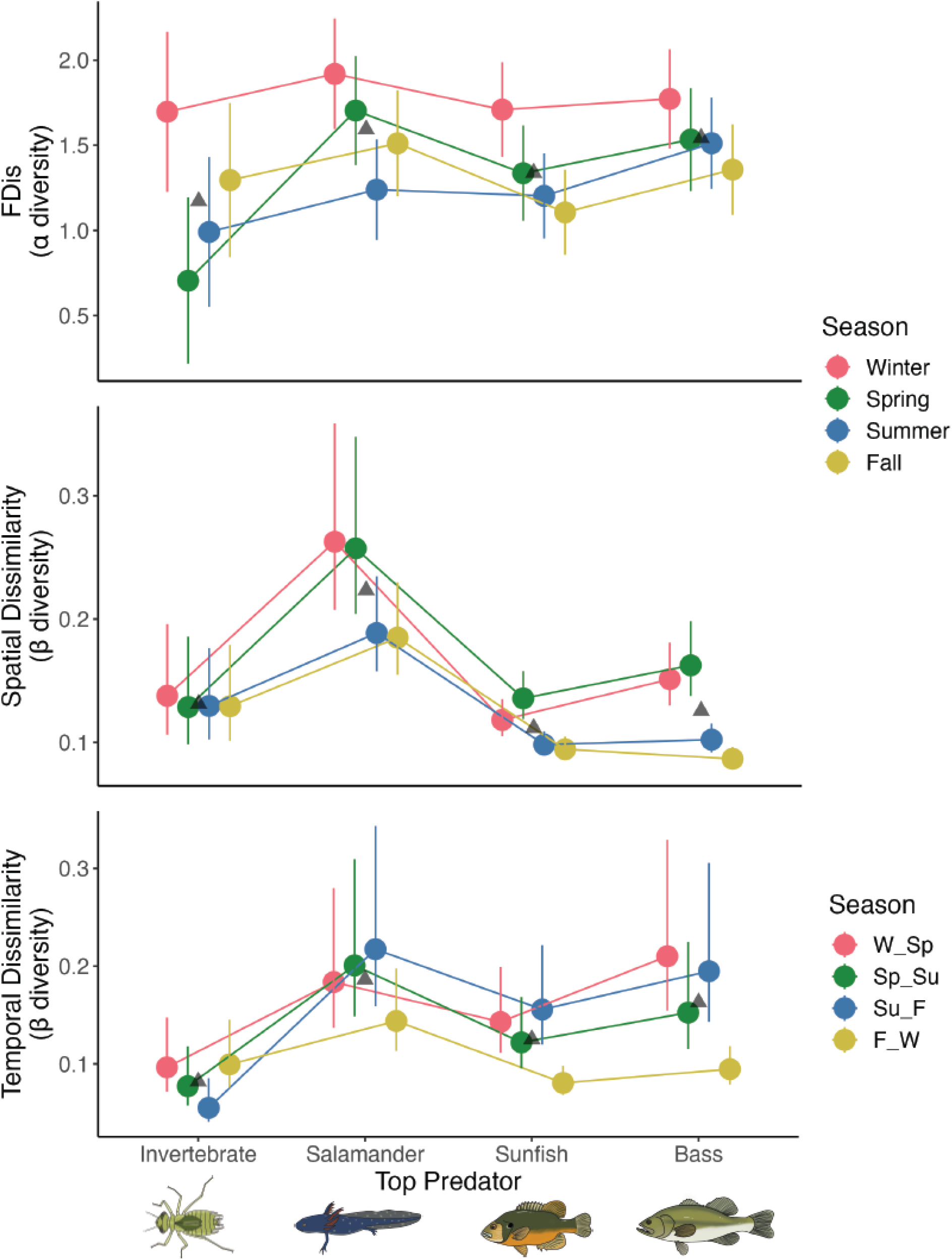
Trait diversity patterns in dragonfly communities across habitats with different top predators. (a) trait α-diversity in ponds (FDis), (b) β diversity indicating differences across ponds within the same top predator, and (c) temporal β diversity indicating temporal turnover of traits across seasons within ponds. Colors indicate season (or season pairs in c) and bars represent 95% confidence intervals.

### Spatial (regional) β- diversity

Top predators generally determined how much trait compositions varied between different pond communities (spatial dissimilarities) (χ^2^ = 21.1139, p = 0.0002), although this effect varied with the year (top predator * year: χ^2^ = 86.277, p < 0.0001) and season (top predator*season: χ^2^ = 56.1658, p < 0.0001). Spatial dissimilarity within seasons was always greatest in salamander ponds (**Fig 3b**), but the greatest differences between salamander ponds and all others occurred in the winter and spring (**Fig 3b**). Sunfish and bass pond dissimilarities were comparable to invertebrate ponds in winter and spring but showed the lowest dissimilarities in summer and fall (**Fig 3b**). In contrast, invertebrate pond dissimilarities were very similar across seasons (**Fig. 4b**). Finally, spatial trait dissimilarity also varied across years (χ^2^ = 10.2170, p = 0.017). The distribution of randomized values simulating equal sample sizes was similar to the observed distribution (**Appendix S1: Fig. S5**).

### Temporal β-diversity

Trait composition within ponds changed with time (temporal dissimilarities) but how much it changed was determined by top predators (χ^2^ = 9.11, p = 0.049) and year (χ^2^ = 8.851, p = 0.004). Salamander ponds showed on average the highest temporal dissimilarity in trait composition and invertebrate ponds the lowest. However, how much temporal diversity differed between trop predator ponds varied with seasonal transitions (predator* season interaction: χ^2^ =23.766, p = 0.005). Temporal dissimilarity between invertebrate vs salamander vs both fish ponds was highest for the spring-summer transition and lowest for fall-winter transition (**Fig. 4c**).

## Discussion

Competition should play a key role in community assembly processes and spatial and temporal biodiversity patterns in natural communities. However, support from natural communities is mixed and previous studies often failed to pinpoint the signals of competition in the biodiversity patterns of natural communities (Münkemüller et al. 2020). This mixed support is not surprising given that competition can shape diversity patterns in various and even opposing ways (e.g. result in low or high functional diversity) depending on the specific underlying mechanism (Mayfield and Levine 2010). Furthermore, it can be masked by other “filtering” processes (e.g. strong abiotic filters) (HilleRisLambers et al. 2012, Kraft et al. 2015). All these factors make it extremely challenging to pinpoint the role of competition and draw a general conclusion from observational data (Münkemüller et al. 2020). Here, we’ve taken a different approach: rather than attempting to indirectly infer competition from diversity patterns, we used a natural competition-predation gradient to examine how competition shapes biodiversity patterns. Overall, results from this “natural experiment” confirm that competition acts as an important deterministic force that shapes biodiversity patterns across local, regional, and temporal scales and provides new insights into the underlying mechanisms.

At the local scale competition can theoretically either increase, decrease, or have no clear effect on the α-diversity of traits or species within communities, depending on what type of competition is dominant (Mayfield and Levine 2010, Münkemüller et al. 2020). We found that trait α- diversity was clearly the lowest within communities that had the weakest predation pressure (invertebrate ponds) compared to intermediate and high predation communities (i.e. salamander and fish ponds) although this effect varied across seasons. This increase in diversity with predation pressure is consistent with empirical and theoretical studies indicating that predation can reduce or even prevent competitive exclusion of species by keeping densities of dominant competitors low (e.g. keystone predation)(Murdoch 1969, Paine 1969, Morin 1981, Holt and Lawton 1994, Leibold 1996, Gurevitch et al. 2000, Saleem et al. 2012). Consistent with this prediction, we also typically found the highest species richness in these ponds (Van Allen et al. 2017), further supporting the hypothesis that predation reduces the probability of competitive exclusion. The decrease in local trait diversity in habitats with high competition indicates that competition between species decreases local trait diversity in our system, suggesting that competition results in the exclusion of species with certain traits.

In contrast to a rich body of work on α-diversity patterns (Münkemüller et al. 2020), surprisingly few empirical or theoretical studies examine trait diversity at larger spatial or temporal scales (β-diversity). Yet, comparing diversity patterns across different scales can help to rule out alternative hypotheses and pinpoint the role of competition in shaping biodiversity. At the regional level, we found that spatial dissimilarity (β-diversity) showed a humped-shaped relationship along the competition-predation gradient: it peaked at an intermediate level of predation (salamander predators), but it was very similar across ponds at either end of the competition-predation gradient. Theory predicts that dissimilarity among communities should decline when deterministic processes such as strong habitat filters increase while the opposite happens when stochastic processes dominate (Chase and Myers 2011). The decline in spatial trait diversity indicates that predators act as strong deterministic filters. This is further supported by the fact that only a small subset of all the observed trait combinations across species are found in ponds with fish, and this subset set of traits is virtually identical for both types of fish predators, and it follows similar seasonal patterns. Our results are consistent with previous studies indicating that certain morphological traits reduce risk of fish predation in odonates (Wellborn et al. 1996, Stoks and De Block 2000, Stoks and McPeek 2006) and are similar to spatial biodiversity patterns observed at the species level. Predation thus acts as a homogenizing, deterministic filter at the regional level for both trait and species diversity.

In contrast to predation, competition has been cited as a factor promoting stochastic processes (Chase 2007, 2010). This perception is based on studies indicating that spatial β- diversity at the species level is typically highest in the absence of predation when competition should be strongest (Chase et al. 2009, Van Allen et al. 2017). However, we found that trait β- diversity was highest at intermediate levels of predation and competition and declined when competition or predation was strongest. This pattern is in contrast to what we observed previously in the same system at the species level and we also showed that both have similar species richness on average (Van Allen et al. 2017). So what is driving this discrepancy at the species and trait level?

Competition is only expected to be a stochastic force that increases β-diversity when priority effects are present. With priority effects, the outcome of competition depends on the relative arrival order (e.g. early excluding late arrivals). If arrival order varies across sites or with sites across time then what species persist and which become excluded should also vary between communities and within communities over time (Chase 2003, Fukami 2015). However, this does not have to be true at the trait level. If species compete for a limited “set of niches”, we expect trait combinations to be still similar across space and time. But given enough functional redundancy among species in the regional pool (i.e. species with similar trait combinations) priority effects would still promote variation in species composition across sites. In this scenario, we expect high β-diversity at the species level but low diversity at the trait level. On the other hand, with limiting similarity, the traits of the first arrivers could determine which of the later arriving species can establish themselves, i.e. those that are too similar to earlier arrivers will be excluded (MacArthur and Levins 1967, Meszéna et al. 2006). In the latter scenario, we expect trait combinations to change across both habitats and time assuming that arrival order naturally varies across time and space. However, if arrival order does not vary and/or priority effects are absent in the system, then species and trait diversity should both be similar across time and space.

The biodiversity patterns in our systems support the “limiting niches” scenario. We found that spatial β-diversity of traits was just as low with high competition as with high predation. Furthermore, the lowest temporal β-diversity was found in habitats with only invertebrate predators, indicating that increasing competition promoted trait convergence across communities. In contrast, temporal β-diversity at the species level was highest in the same type of habitat. Together, these results indicate that competition selects for a relatively fixed combination of traits, while priority effects and functional redundancy across species could still promote variation in species composition. Overall, our study revealed that competition can act as a strong deterministic habitat filter at the trait level, but priority effects can still promote stochasticity at the species level given functional redundancy in the regional species pool.

The deviation of several of our findings from past studies on species diversity (Van Allen et al. 2017) provides further evidence that ecological processes cannot easily be inferred from patterns in species diversity (Münkemüller et al. 2020). For instance, in habitats with strong predation trait combinations did not change dramatically across seasons like species composition, but remained surprisingly constant. Likewise, high-competition habitats were not more strongly influenced by stochastic processes than high-predation habitats as is assumed from studies on species diversity. Consistent with previous studies (McGill et al. 2006, Baraloto et al. 2012, Purschke et al. 2013, Funk et al. 2017, Münkemüller et al. 2020), these results indicate that functional diversity is often a more informative metric than species diversity in ecological studies and thus should be included when possible.

## Conclusions

Inferring processes that drive biodiversity patterns and community assembly is always challenging. The required scale is typically too large for experiments and observational studies typically cannot provide a final answer about the mechanisms of community assembly. Here we show that we can still use observational data to help identify underlying processes by using natural gradients (or experiments) and examine patterns at different spatial and temporal scales and test theoretical predictions. Similar approaches in other systems thus provide a fruitful venue to gain a deeper understanding of the factors that shape biodiversity patterns. Our results also revealed that competition can act as both a strong deterministic driver at the trait level and as a stochastic driver at the species level. These results emphasize the importance of combining both types of data to help identify how important different processes like niche sorting and priority effects are in a given system and maintaining biodiversity in natural communities.

## Acknowledgements

We thank the wildlife biologists of the U.S.D.A. Forest Service Wildlife and Silviculture Southern Research laboratory, particularly D. Saenz, for locating the study ponds and conducting preliminary surveys of the predator communities. We also thank A. Roman and G. Ross for help collecting samples and P. Delclos for help with keying odonates. This work was supported by NSF DEB-1256860, DEB-0841686 and DEB- 1655626 to V.H.W. Rudolf.

## Conflict of interest

The authors declare no conflict of interests

## Notes

### Competing Interest Statement

The authors have declared no competing interest.

