## Supplemental material for "Predation and competition drive trait diversity across space and time"

**Appendix S1 for:**

**Predation and competition drive trait diversity across space and time**

**Zoey Neale & Volker H. W. Rudolf***

Graduate Program in Ecology & Evolutionary Biology, BioSciences

Rice University,

Houston, TX 77005, USA

Includes supplemental **Table S1** and **Figures S1-S5**

**Table S1** Odonate species included in this study.

| Suborder | Family | Genus | Species |
| --- | --- | --- | --- |
| Anisoptera | Aeshnidae | *Anax* | *junius* |
| Anisoptera | Aeshnidae | *Anax* | *longipes* |
| Anisoptera | Aeshnidae | *Basiaeschna* | *janata* |
| Anisoptera | Aeshnidae | *Epiaeschna* | *heros* |
| Anisoptera | Corduliidae | *Epitheca* | *cynosura* |
| Anisoptera | Corduliidae | *Epitheca* | *semiaquea* |
| Anisoptera | Libellulidae | *Celithemis* | *fasciata* |
| Anisoptera | Libellulidae | *Celithemis* | *ornata/verna* |
| Anisoptera | Libellulidae | *Erythemis* | *simplicicollis* |
| Anisoptera | Libellulidae | *Erythrodiplax* | *minuscula* |
| Anisoptera | Libellulidae | *Ladona* | *deplanata* |
| Anisoptera | Libellulidae | *Libellula* | *spp.* |
| Anisoptera | Libellulidae | *Pachydiplax* | *longipennis* |
| Anisoptera | Libellulidae | *Pantala* | *flavescens* |
| Anisoptera | Libellulidae | *Perithemis* | *tenera* |
| Anisoptera | Libellulidae | *Plathemis* | *lydia* |
| Anisoptera | Libellulidae | *Tramea* | *carolina* |
| Zygoptera | Coenagrionidae | *Enallagma* | *aspersum* |
| Zygoptera | Coenagrionidae | *Enallagma* | *divigans* |
| Zygoptera | Coenagrionidae | *Enallagma* | *dubium* |
| Zygoptera | Coenagrionidae | *Enallagma* | *geminatum* |
| Zygoptera | Coenagrionidae | *Enallagma* | *signatum* |
| Zygoptera | Coenagrionidae | *Enallagma* | *vespersum* |
| Zygoptera | Coenagrionidae | *Ischnura* | *posita* |
| Zygoptera | Coenagrionidae | *Nehalennia* | *integricollis* |
| Zygoptera | Coenagrionidae | *Telebasis* | *byersi* |
| Zygoptera | Lestidae | *Lestes* | *australis* |

**Figure S1** Mean trait values (see table S1 for trait descriptions) of odonates within pond predator types across years in winter.

**Figure S2** Mean trait values (see table S1 for trait descriptions) of odonates within pond predator types across years in spring.

**Figure S3** Mean trait values (see table S1 for trait descriptions) of odonates within pond predator types across years in summer.

**Figure S3** Mean trait values (see table S1 for trait descriptions) of odonates within pond predator types across years in fall.

**Figure S4** Distributions of mean FDis (a), spatial dissimilarity (b), and temporal dissimilarity (c) for each of 500 random draws of three ponds of each predator group.

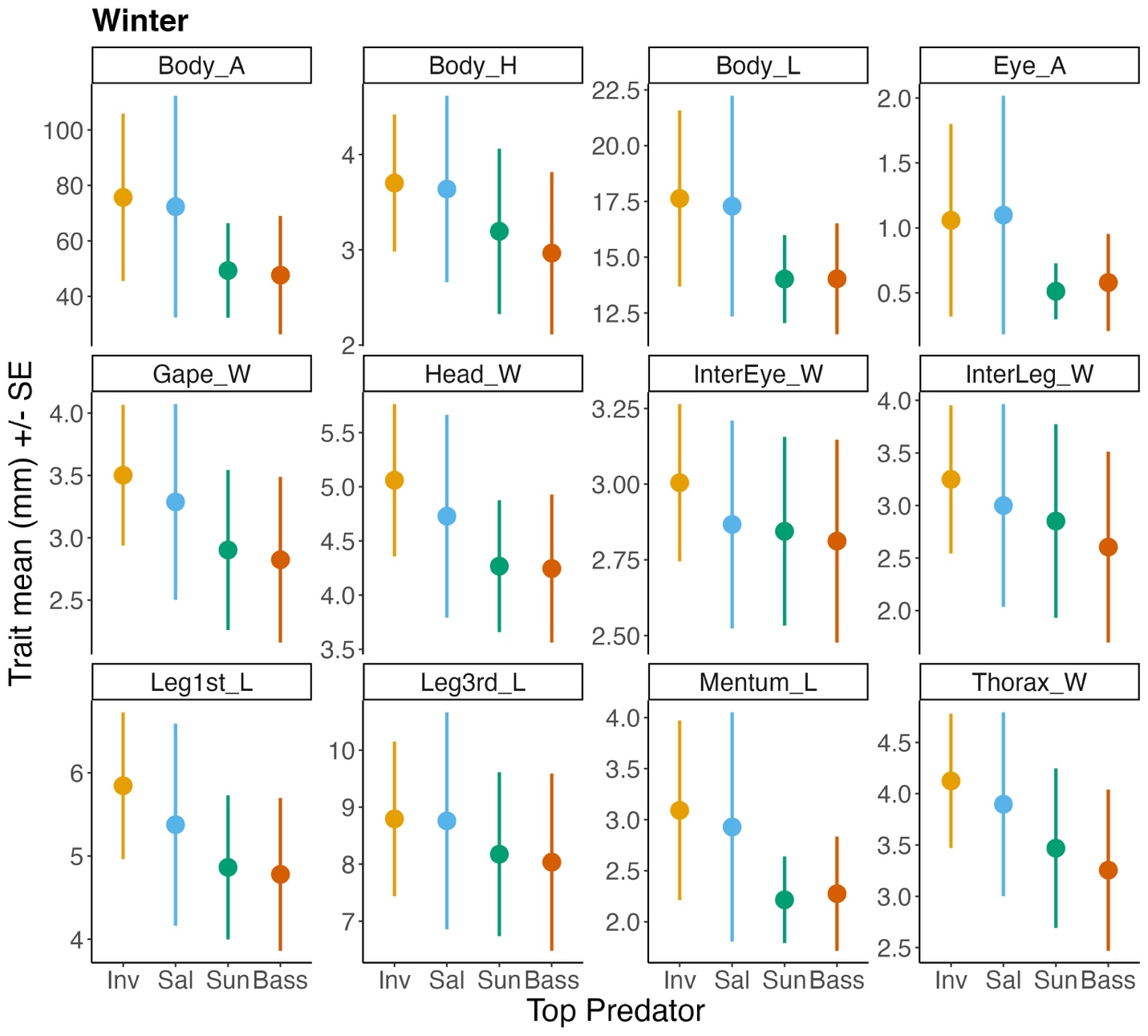

**Figure S1**

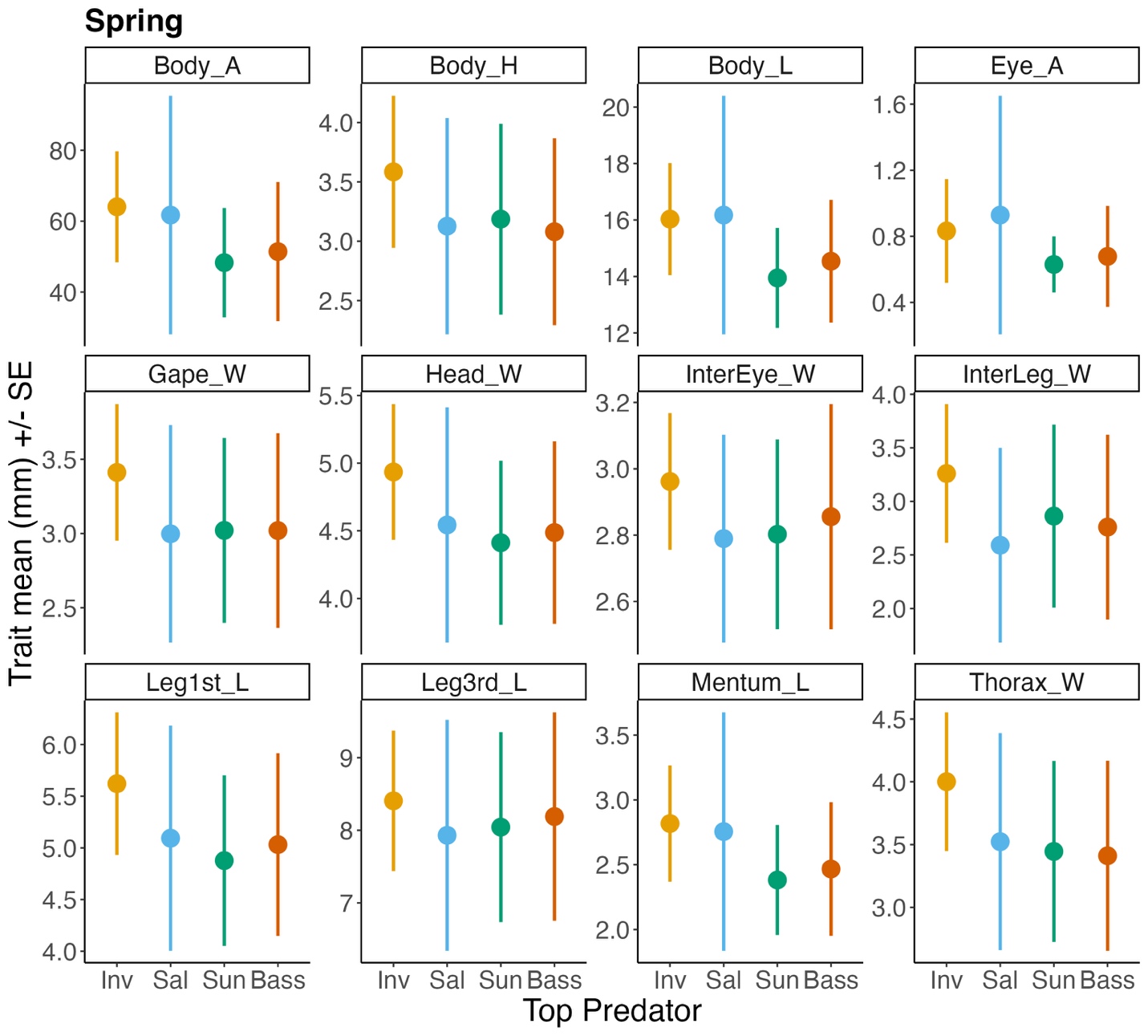

**Figure S2**

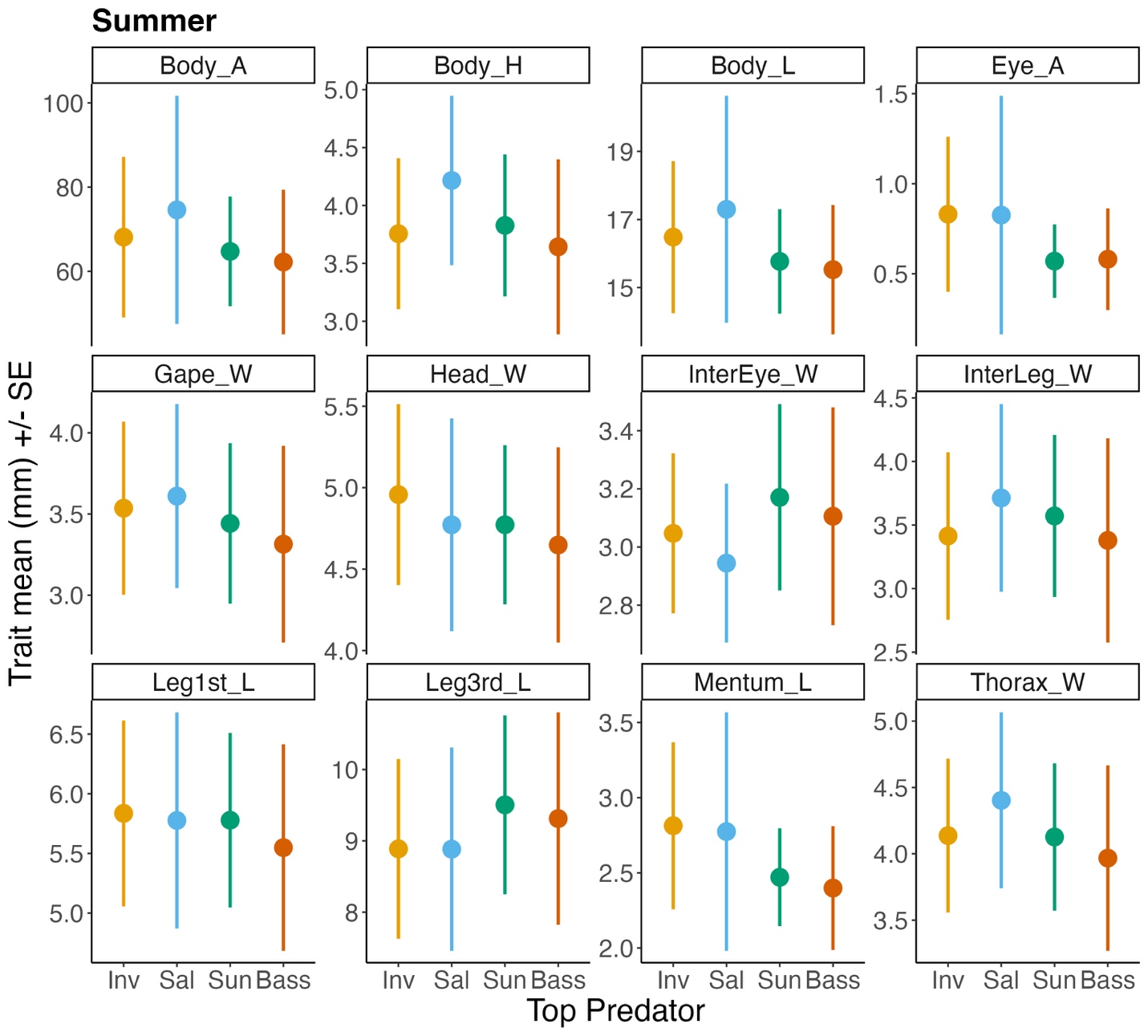

**Figure S4**

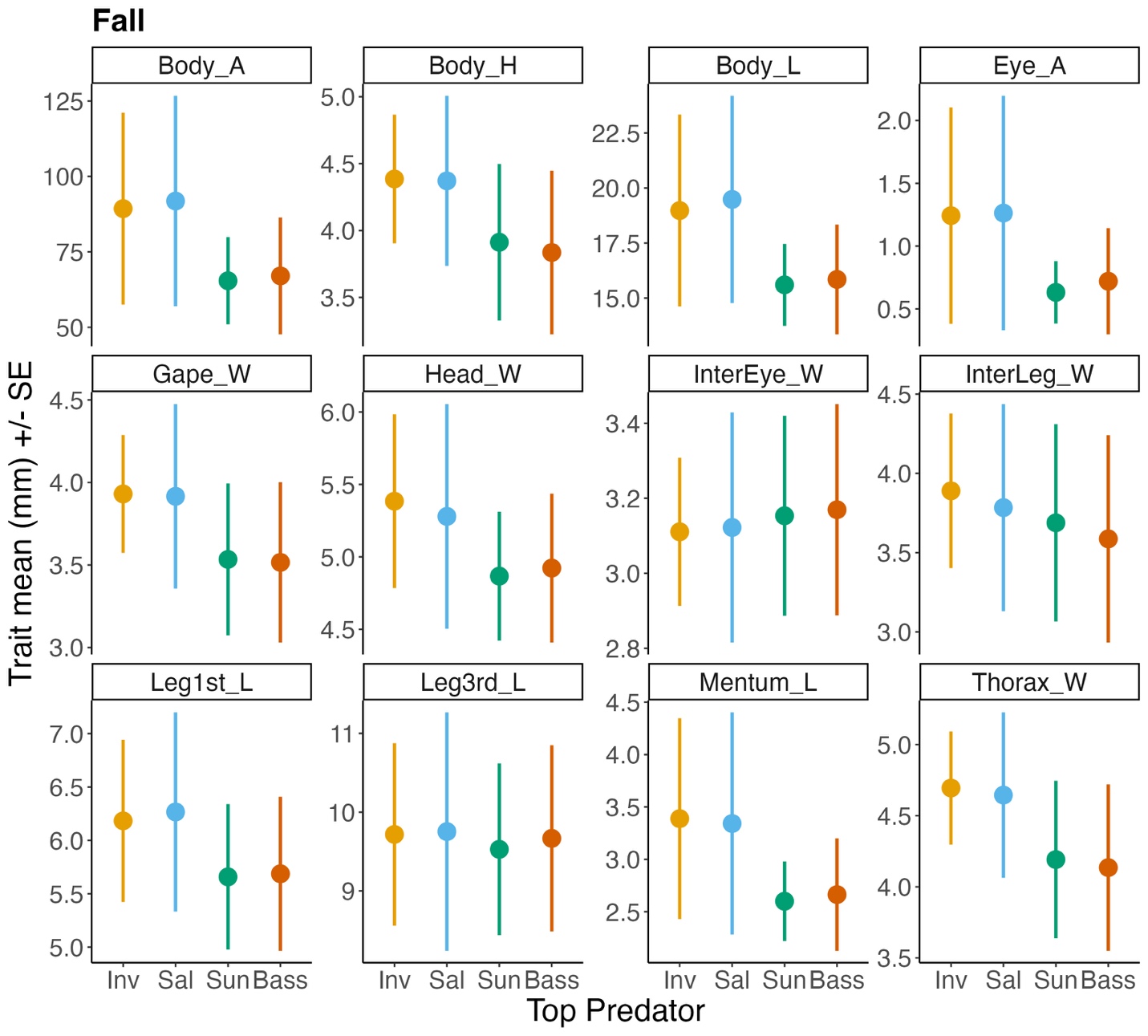

**Figure S4**

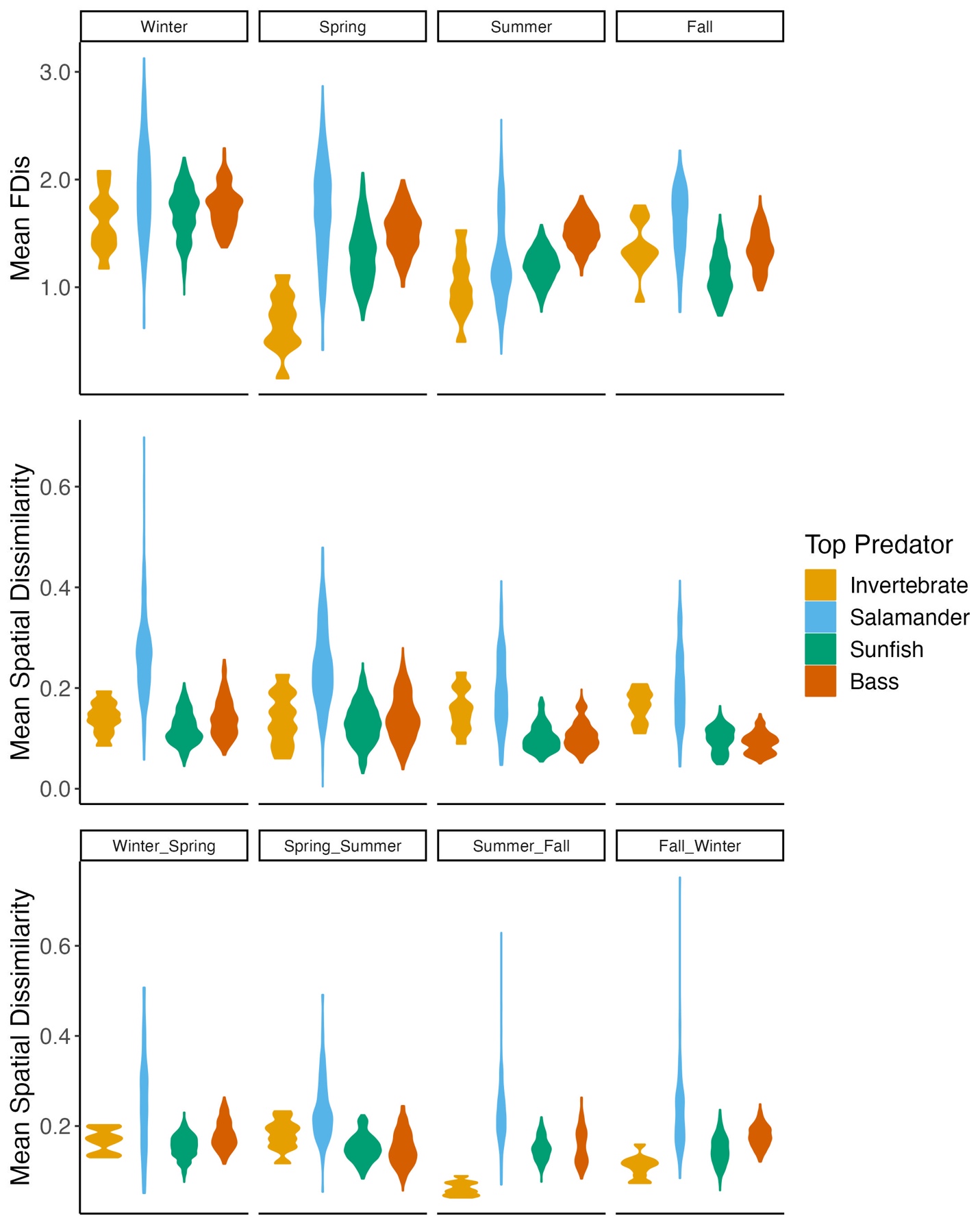

c)

b)

a)

**Figure S5**
